# Novel hypervariable erythrocyte surface expressed recombinant proteins show promise as serological markers of exposure to *Plasmodium falciparum* infection

**DOI:** 10.1101/2023.10.31.564916

**Authors:** Hristina Vasileva, Ana Chopo-Pizarro, Michael Ooko, Elin Dumont, Lindsey Wu, Isaac Ssewanyana, Anna Last, Kevin K.A. Tetteh

## Abstract

Malaria caused by *Plasmodium spp*. leads to significant morbidity and mortality, particularly in Sub-Saharan African countries with *Plasmodium falciparum* being the predominant infectious species. The pathogenesis of *P. falciparum* depends on multiple host and parasitic factors, one of which is the evasion of host immune response due to antigenic variability during the blood stage of infection. The understudied infected erythrocyte expressed protein families STEVOR and RIFIN characterize with antigenic hypervariability and are associated with clinical outcome of the infection and protective acquired immunity based on their topology and localization. We have used two molecular tag methods for successful expression of members of STEVOR and RIFIN protein families as recombinant proteins in *E. coli* expression system. We have further established the antigenicity of those recombinants and have used Ugandan cohort samples with various *P. falciparum* infectious status and have compared the seropositivity rate to those recombinants in different age groups against already established short- and long-term markers of infection. We have demonstrated age-dependent immunity acquisition against the tested recombinants, and we have suggested the potential use of STEVOR and RIFIN recombinants as novel markers of *P. falciparum* infection in serosurveillance. Due to the hypervariability of those protein members we propose that further, more extensive research using a library of expressed variants is needed to strengthen the conclusions made in this study.

## 1. Introduction

Malaria is a vector-borne disease caused by parasites from the genus *Plasmodium* and transmitted by *Anopheles* mosquito species, responsible for approximately 619,000 deaths and 247 million infectious cases in 2021, with the greatest burden of disease in sub-Saharan Africa ^1,2^. There are five main parasite species which cause human malaria, including a zoonotic form, of which *Plasmodium falciparum* (*Pf*) is the most virulent, responsible for over 99% of malaria cases, the majority of which occur in children under five years of age and pregnant women ^1,3^. Despite the observed reduction of malaria cases between 2000 and 2015 due to malaria control strategies, cases have been increasing significantly since 2016 due to several factors such as the rise of drug-resistant parasites and vector insecticide resistance, coupled with issues of regional political instability ^1,4,5,6^. More recently, the WHO estimated that interruptions to malaria control efforts and case management caused by the SARS-CoV-2 pandemic resulted in an additional 13.4 million cases ^1^.

Pathogenesis of malaria depends on multiple parasite and host factors, causing different clinical outcomes and disease severity ^7^. Some of the parasite’ s virulence is due to evasion of the human host immune system during the blood stage of the infection. *Pf* infected erythrocytes (IE) adhere to the epithelial wall of blood vessels, via cytoadherence, preventing the clearance of the IE from the blood stream. The formation of rosettes around IE ensures more efficient spread of infection ^7^. Sequestration and cytoadherence are characteristic *Pf* virulence factors, enabled by parasite derived proteins expressed on the surface of IEs. These proteins possess antigenic properties and antibodies produced following exposure are associated with acquired protective immunity to *Pf* malaria ^8^. Moreover, IE surface expressed antigens are associated with antigenic variability, termed Variant Surface Antigens (VSA) ^8^. These antigens are the protein products of multi-copy gene families, grouped into several protein families: *P. falciparum* erythrocyte membrane protein 1 (*Pf*EMP1), repetitive interspersed protein family (RIFIN), sub-telomeric variable open reading frame family (STEVOR), *P. falciparum* Maurer’ s cleft two transmembrane domain family (*Pf*MC-2TM) and surface-associated interspersed protein family (SURFIN) ^9^. These proteins are translocated from the blood stage *Pf* parasites (merozoites and schizonts) to the surface of the IE via protein trafficking through the Maurer’ s cleft, a parasite derived membranous structure, and are expressed on the IE membrane, protruding into the extracellular space ^9^. Thus, they can be regarded as potential novel therapeutic targets, as has been investigated in previous studies on *Pf*EMP1 ^10^.

RIFIN and STEVOR protein families are encoded by approximately 180 *rif* and 40 *stevor* gene copies per parasite, respectively, where one gene from each family is expressed at a time per individual parasite, following an infection stage depended successive expression ^9,11^. Members of each family differ mostly in their hypervariable region, believed to be the only domain exposed to the circulation that possesses antigenic epitopes, until a study using peptide microarrays showed that in *Pf* infected populations, there were similar levels of serorecognition and seroreactivity to both the semi-conserved (SC) and the large hypervariable (V2) protein domains of STEVORs and RIFINs ^12,13^. Moreover, both SC and V2 domains were found to be associated with *Pf* exposure and potentially with clinical outcomes of the infection ^12^. Seroreactivity and serorecognition to both protein families have been demonstrated to be age and exposure dependent, with high reactivity in adults compared to children and higher domain recognition in individuals with clinical malaria as opposed to sub symptomatic infections ^12^.

This is the first study demonstrating successful expression of isolated domains of members of the STEVOR and RIFIN protein families as recombinant proteins. Antibody reactivity to previously established markers of infection have been used as juxtaposition to classify the nature of antigenicity of these new targets^14,15,16^. This study aims to fill in the gaps in *Plasmodium falciparum* proteome knowledge by investigating possible new markers for measuring exposure to infection.

## 2. Materials and Methods

### 2.1 Ethics

This study has received ethical approval by the Ethics Committee of the London School of Hygiene and Tropical Medicine (LSHTM) (reference number: 21505). The serum samples used in this study come from the Program for Resistance, Immunology, Surveillance, and Modelling of Malaria in Uganda longitudinal cohort (PRISM), approved by the Ethics Committee of LSHTM (reference number: 15823) and the Research and Ethics Committee of the Makerere University School of Medicine in Kampala, Uganda (reference number: 2011-167), and the Mapping Malaria and Neglected Tropical Diseases on the Bijagos Archipelago of Guinea Bissau (DTNMaPa), approved by the Ethics committee of LSHTM (reference number: 22899) and the Comite Nacional de Eticas de Saude (CNES) in Bissau, Guinea Bissau (reference number: 076/CNES/INASA/2017).

### 2.2 Design and expression of STEVOR and RIFIN recombinant proteins

Six STEVOR (PF3D7_1300900, PF3D7_1254100, PF3D7_0115400, PF3D7_0300400, PF3D7_0832000, PF3D7_0832600) and three RIFIN (PF3D7_0833200, PF3D7_1041100, PF3D7_0732400) protein sequences constructed from the 3D7 *Plasmodium falciparum* reference strain were initially selected^12^. Protein sequences were downloaded from the PlasmoDB database (Plasmodium Genomic Database, RRID:SCR_013331) ^17^ and multiple alignment of the sequences was performed for each protein family using Clustal2X ^18^. Additional inspection of the alignments was conducted manually to identify misalignments. Both protein families present a similar structure containing a signal peptide (SP), a semiconserved domain (SC), genetically different between the families conserved domain (C), two hypervariable domains (V1 and V2) and 2 transmembrane domains (2TM), as well as a translocator signaling element (PEXEL motif) (Figure 1A) ^19^. The SC and V2 domains of the amino acid sequences were isolated for both the STEVORs and RIFINs, using published data on domain architecture literature and a transmembrane domain prediction server (TMHMM v.2.0) ^20^. The edited sequences were then realigned using the ‘msa’ package on R computational platform (v3.6.3; R Core team 2020) ^21^. The V2 and SC portions of three STEVORs (PF3D7_1300900, PF3D7_0832000 and PF3D7_0832600) and one RIFIN (PF3D7_0732400) proteins were selected as representative sequences, based on divergence using protein sequence phylogenetic trees (IQtree) ^22^.

**Figure 1:**
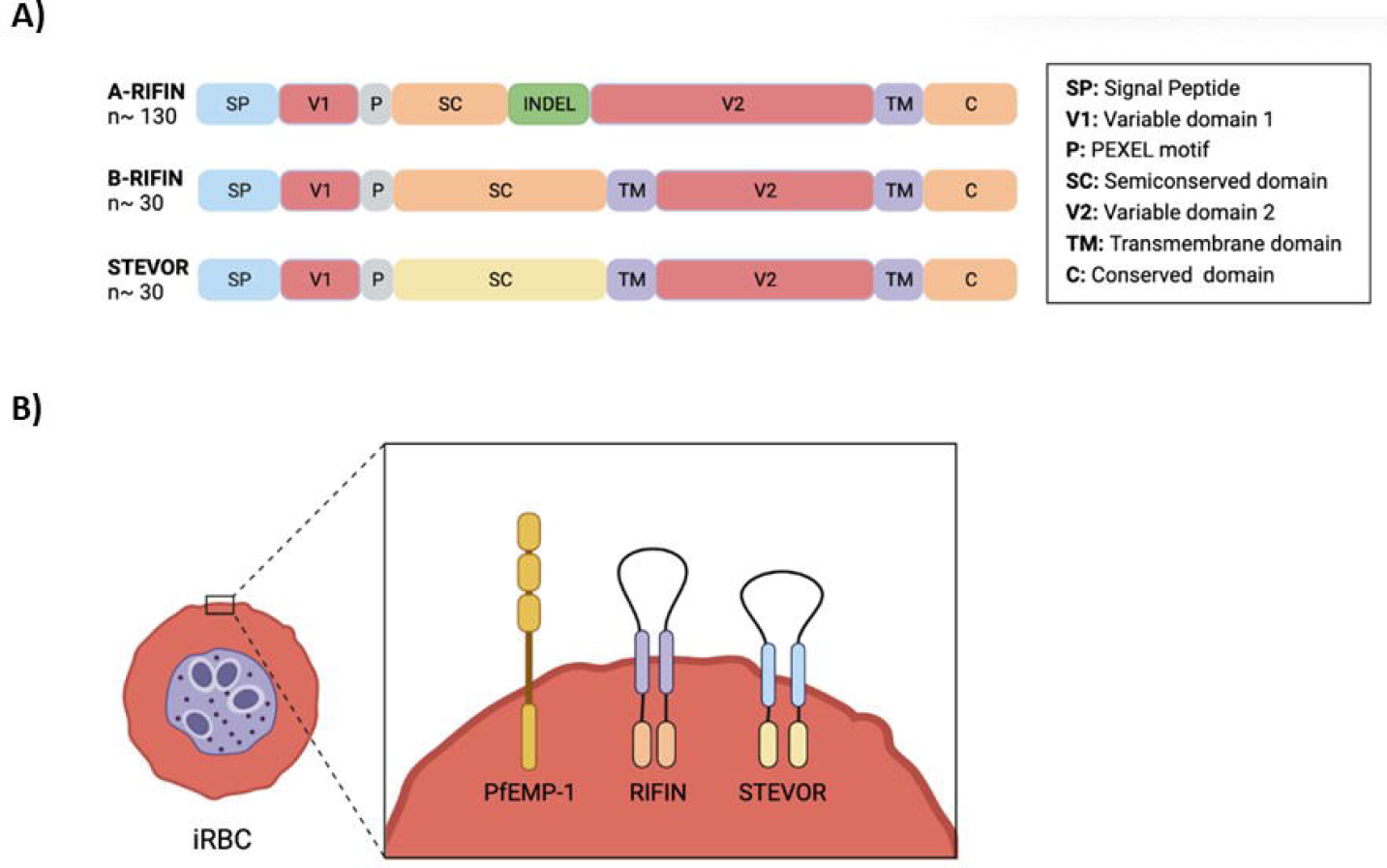
(A) General structure of the RIFIN (A and B) and STEVOR proteins showing the domain architecture of the proteins (B) and their predicted location at the surface of infected red blood cells in relation to *Pf*EMP1. Loops protruding outside the cell membrane correspond to the large variable domain (V2), flanked by the transmembrane domains (shown in lilac for the RIFIN and blue for the STEVOR proteins). Transmembrane regions were identified using the TMHMM v.2.0 transmembrane domain prediction server. (Figure adapted by Zhoe E. Albert *et. al* (2019), *mSphere;* Created with BioRender.com)

The selected SC and V2 domains were expressed as recombinant antigens in *E. coli*, as described elsewhere ^23,24^. Briefly, the SC and V2 domains were cloned into the pET15b expression vector as dual His (N-terminal) and Myc (C-terminal) tagged recombinant constructs (Biomatik, USA). A duplicate set of expression constructs were also cloned into the pGEX-4T-1 expression vector as single-tagged GST (N-terminal) recombinant constructs (Biomatik, USA), resulting in a total of fourteen recombinant proteins. BL21(DE3) chemically competent *E. coli* (Trans, China) was transformed with either the pET-15b or pGEX-4T-1 plasmid constructs, each containing an Ampicillin (Amp) resistance cassette. The transformed bacteria colonies were cultured in ZY auto-induction media supplemented with100 μg/ml Amp at 37°C, 150 rpm overnight (∼16 hr) ^24^. The cells were pelleted at 7500 x g, resuspended in 1xPBS and lysed using an LM20 microfluidizer (Analytik LTD, UK) under 18,000 psi pressure units. The dual His-Myc tagged recombinant proteins were affinity purified using an ÄKTA Pure purification system (Cytiva, USA) nickel chromatography. Bound proteins were eluted using an imidazole gradient. The eluted His-Myc tagged recombinants were further concentrated using 3kDa ultra concentration units (Merck, Germany). The GST-tagged recombinant proteins were affinity purified using Glutathione Sepharose 4B beads and purified in batch mode. No further concentration of the GST tagged proteins was required. Both sets of recombinant proteins were quantified using the Bradford protein quantification assay (Bio-Rad, UK).

### 2.3 Serum pools

An ELISA assay was developed to evaluate the reactogenicity of the STEVOR and RIFIN recombinant proteins ^25^. A two-fold dilution series was prepared for each recombinant protein in coating buffer (1.59g/l Na_2_CO_3_ and 2.93g/l NaHCO_3_) starting at 4 μg/ml, down to 0.5 μg/ml. The dilution series for each recombinant was assayed against 6 positive control serum pools in addition to a malaria naïve negative control pool (PHE: Public Health England)^26^. All sera were assayed at a 1/200 dilution. Following incubation, plates were developed using 3,3’, 5,5’ -tetramethylbenzidine (TMB; Tebubio: #TMBW-1000-01) and read at 450 nm. Generating a specific serological control for this study was imperative since it was practically unknown which variants of the STEVOR and RIFIN families were serorecognised by the tested population, as well as the control pooled populations. The control serum pools tested for control selection were as follows. Two in-house control pools together with an international reference standard were: *P. falciparum* hyperimmune serum pool based on Tanzanian adults positive for *Pf* infection (CP3), Gambian adults hyperimmune control pool (Brefet) and the 10/198 WHO 1^st^ international *Pf* serum standard based on Kenyan adult samples (WHO)^27^. Three additional pooled controls added to the panel were: a pool of 40 Ugandan serum samples from individuals aged above 18 years with confirmed infection using Loop-mediated isothermal amplification (LAMP) and confirmed clinical disease (PRISM1), the highest quartile of 80 tested individuals from Uganda of variable ages with calculated percentile antibody reactivity against in-house established markers of *Pf* infection (PRISM2), and the highest quartile of 40 tested individuals from the Bijagos Archipelago, Guinea-Bissau of above 18 years with confirmed infection according to *Pf* 18S qPCR, with calculated percentile antibody reactivity against already tested in-house established markers of *Pf* infection (DTNMaPa). Highest population quartiles of antibody reactivity for PRISM2 and DTNMaPa were calculated based on median fluorescence intensity (MFI) data of antibodies against the recombinant proteins *Pf*AMA1, *Pf*MSP1.19, GLURP.R2, Etramp5.Ag1 and HSP40, obtained from MagPix multiplex bead-based array^28^ (Vasileva *et. al;* DTNMaPa unpublished data).

### 2.4 Detection of optimum recombinant protein antigenicity

After confirming protein seroreactivity using ELISA, the recombinant proteins were chemically coupled to MagPlex microsphere beads (Luminex) using an 8-fold 6-point dilution series according to established methods^29^. Briefly, titration of the coupled beads was performed against 5-point 2-fold dilutions of two positive control pools down selected from the ELISA data (CP3 and PRISM1) and a negative (PHE) control serum pool starting from 1/200 down to 1/1600 serum dilutions. MFI data was used to calculate the 50% maximum effect of antibody titers, also known as the EC50 point on a sigmoidal curve. Four parameter logistic regression was used to calculate the EC50 point per dilution curve, and the median EC50 point across all dilutions of both controls for each recombinant was selected as an optimum bead coupling concentration.

### 2.5 Sample selection

Plasma samples used for this study were collected during the first series of the cohort studies from the Program for Resistance, Immunology, Surveillance and Modelling of Malaria in Uganda (PRISM), conducted between October 2011 and June 2016 in three areas of Uganda: Kihihi, Kanungu District; Walukuba, Jinja District; and Nagongera, Tororo District.^30^ A total of 505 samples were selected from the 2013 pre-intervention (n=310), and 2016 post insecticide treated nets distribution (ITN) and indoor residual spraying (IRS) interventions (n=195), from the Tororo district. Tororo is a rural setting located in the southeastern region of Uganda near the border with Kenya, characterized with high malaria transmission intensity ^31^. Selected samples were collected from April until the end of June for both timepoints, to include the highest peak of annual malaria transmission, indicated in the sample distribution plot in Supplementary Figure 2. Study participants were closely monitored to identify malaria cases via passive case detection (participants seeking health care when feeling ill) as part of the longitudinal active surveillance (routine visits every 90 days), with cases identified by microscopy and/or LAMP^28^.

### 2.6 Multiplex bead-based serology assay

MagPlex magnetic beads were coupled with the STEVOR and RIFIN recombinant proteins, with one expression system per recombinant selected, summarized in Table 1. The recombinants were complemented in the Luminex assay with additional markers of seroprevalence, summarized in Supplementary Table 1 ^32,26,16^. Serum samples were diluted at 1/400 in antibody elution buffer and incubated overnight ^29^. The Luminex serology assay was performed following an optimized standard operating procedure, as described previously ^29^. Briefly, serum samples were incubated with the antigen coupled beads, washed with 1xPBS/T, then incubated with Goat anti-human IgG R-phycoerythrin (R-PE) labelled antibody (Jackson ImmunoResearch©) for signal detection. A six point 5-fold dilution series, starting from 1/10 down to 1/31250 for CP3 and PRISM1 positive controls, was run per plate for obtaining standard curve values, and further used for plate-to-plate variation quality control. Additional controls included two wells of PHE malaria naïve negative controls diluted 1/400 and two wells of background control containing only antibody elution buffer. The plates were read using a MagPix© bioanalyzer and data was obtained in the form of MFI, a proxy measure of antibody titers. MFI data was background adjusted and data quality checked by comparing the control standard curves and using Levy Jennings plots of mean MFI data for assessing plate-to-plate variations. Any plate with data falling outside of the accepted variation of mean plus/minus 3 standard deviations was repeated ^29^.

**Table 1:** Summary of recombinant STEVOR and RIFIN proteins selected. Protein titration plots for the selection of optimum protein concentration (EC50 point of saturation) are displayed as Supplementary Figure 1.

| Reference Name | Recombinant Name* | Molecular Tag | Mol. weight (kDa) | Concentration (µg/mL) | EC50 (µg/mL) |
| --- | --- | --- | --- | --- | --- |
| Pf3D7_1300900 | STEVOR1_SC | GST | 38.02 | 593.30 | 3.70 |
| Pf3D7_0832600 | STEVOR6_SC | GST | 38.55 | 1049.20 | 12.85 |
| Pf3D7_0353200 | STEVOR5_SC | GST | 38.71 | 1109.10 | 2.95 |
| Pf3D7_0732400 | RIFIN3_V2 | GST | 40.26 | 1193.30 | 7.05 |
| Pf3D7_1300900 | STEVOR1_V2 | His-Myc | 4.44 | 842.30 | 3.30 |
| Pf3D7_0832600 | STEVOR6_V2 | His-Myc | 4.66 | 448.20 | 2.90 |
| Pf3D7_0732400 | RIFIN3_SC | His-Myc | 4.90 | 504.30 | 14.00 |

\*Semi-conserved domain recombinants (SC); Large variable domain recombinants (V2)

### 2.7 Data analysis

Antigen-specific seropositivity was calculated using mean MFI data plus three times the standard deviation of the PHE negative serum control per antigen. Samples above the threshold are termed seropositive and percentage seropositivity per antigen per age group was displayed for both sample time points, 2013 and 2016 as a figure using R version 4.2.3 statistical software. The three samples age groups selected were as follow: “6 months – 5 years”; “5 years - 11 years” and “above 18 years”. The proportion percentage seropositivity, accounting for the differences in samples size between the two time points, was used for calculating the odds ratio (OR) of being seropositive in 2016 compared to 2013, briefly as follows. Logistic regression for binomial data distribution model was built to calculate the log odds of being seropositive in 2016 over being seropositive in 2013, calculated using year and age as covariate, including the variables age and year interaction. The logistic regression was adjusted for clustering at the individual level, using robust standard error method, to account for repeated samples from the same individuals across multiple years. Data was exponentiated to obtain OR and confidence intervals (CI). The logistic regression model was built using “glm” function from the “stats” package and displayed as a forest plot using “forestplot” package in R version 4.2.3 statistical software.

## 3. Results

### 3.1 Design and expression of STEVOR and RIFIN recombinant proteins

Dissecting the SC and V2 domains (Figure 1A) was critical for successful expression of the recombinants in *E. coli* system, as incorporating malarial proteins specific domains such as signal peptides and transmembrane domains was likely going to result in protein aggregation into bacterial inclusion bodies, resulting in insoluble and dysfunctional recombinants ^33^. All targets were expressed using both types of plasmids (dual His-Myc tag and single GST tag). For each of the proteins, one type of recombinant was selected based on multiple factors such as volume of successful protein expression, concentration of protein after purification and seroreactivity according to titrations against the controls, summarized in Table 1. GST tagged recombinants showed higher reactivity compared to the His-Myc tagged recombinants (shown in the titration plots in Supplementary Figure 1). The nature of the recombinants could explain this in terms of semi-conserved region versus large variable domain, where the SC is expected to be more serorecognised as this region is more conserved between variants, thus higher cross reactivity is expected. Additionally, to correct for any potential reactivity to the GTS tag itself, MFI values against the GST tag alone per sample could be subtracted from the total sample MFI data. Although this was not necessary as the seropositivity analysis to GST showed that none of the samples have MFI values above the seropositive threshold (shown on Supplementary Table 2). Furthermore, the His-Myc dual tag recombinants show some level of precipitation after protein purification, which is nullified in the GST-tag expression system since the GST molecule acts as a solubility factor ^**34**^. Despite the molecular differences of their weigh, 4.5-8.8 kDa for His-Myc and 31.7-40.3 kDa for GST, both tag systems were comparable in terms of recombinant concentrations, and it did not explain any differences observed in the results, showing that the expression tags did not have a significant effect on the proteins being investigated.PRISM1 is a pool of 40 LAMP positive Ugandan samples that had the highest reactivity against all recombinants, as shown in Figure 2. Moreover, the tested samples in this study were from the same geographical population as the selected control pool, making it the best control candidate for the study. As per the recombinants, there is an overall higher reactivity to the SC domains compared to the V2 domains, more prominently observed in STEVOR1 and RIFIN3, shown in the last plot in Figure 2.

**Figure 2:**
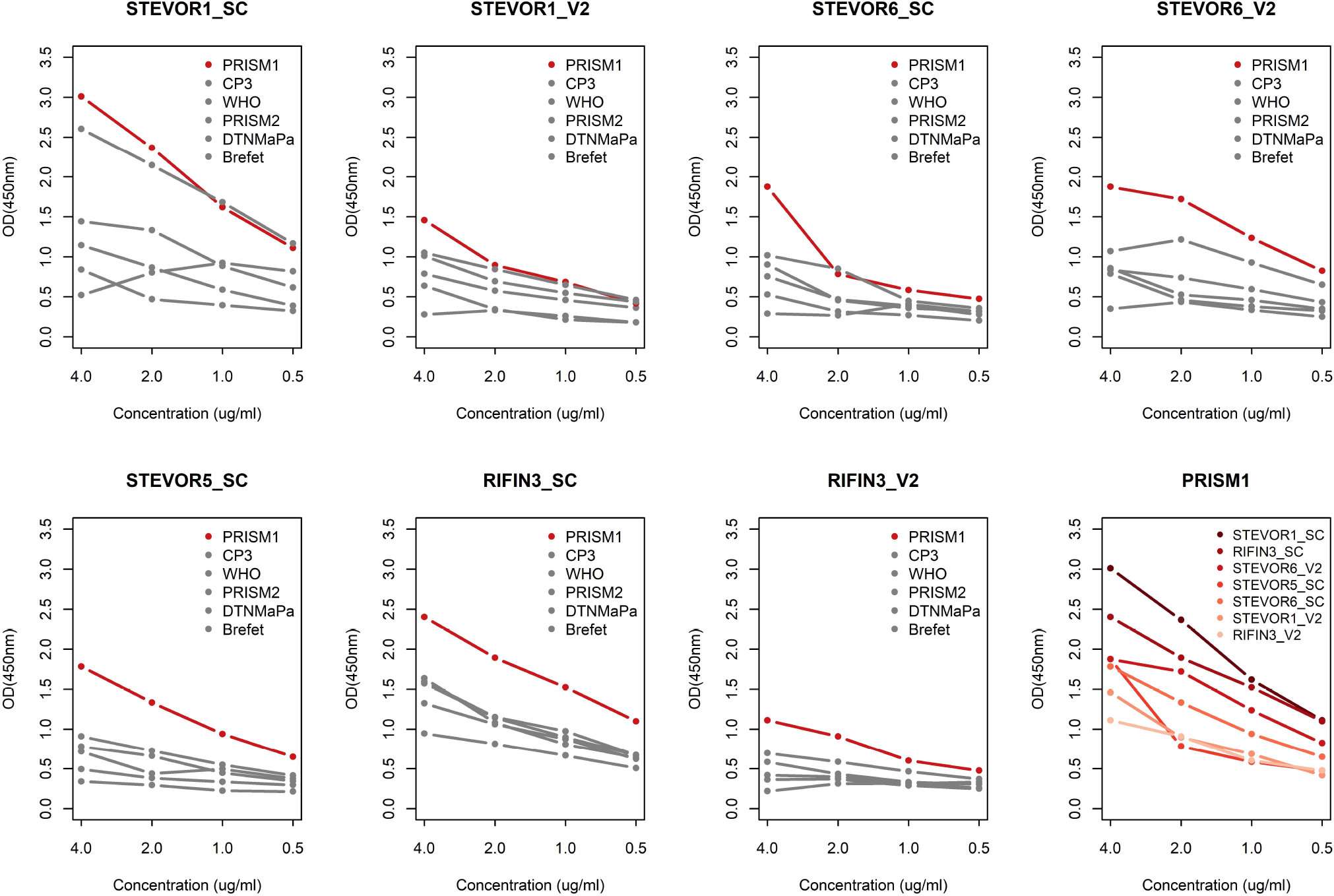
ELISA results of control serum pools (1/200 serum dilution), challenged against 4 steps 6-fold dilution series of each recombinant protein. Brefet: Serum pool of hyperimmune adults from The Gambia; CP3: LSHTM in house serum pool of hyperimmune Tanzanian adults; DTNMaPa: Serum pool of 40 *Pf* infection confirmed adults, with high antibody responses to *Pf* markers of exposure from Bijagos Archipelago, Guinea-Bissau; PRISM1: Serum pool of 40 Ugandan adults with confirmed *Pf* infection and clinical disease; PRIMS2: Serum pool of 80 Ugandan all age individuals with high antibody responses to established *Pf* markers of exposure; WHO: WHO 1^st^ international *Pf* serum from Kenyan adult population. The control serum selected from this experiment is PRISMS1, indicated in red. The latest plot of the series demonstrates the seroreactivity of all recombinants against PRISM1, with color-coded intensity in descending order.

### 3.2 Sample summary statistics

There was a decrease of 37% of samples used in the study between 2013 and 2016 from 310 to 195, respectively, due to a change of the regime in sampling from all samples collected at two time points per year in 2013, to collection of samples over a period of 3 months in 2016 form the same study participants, indicated in Supplementary Figure 2. The decrease in malaria cases confirmed using microscopy and LAMP from 217 to 40 cases, accounting for more than 50% decrease, summarized in Table 2, is explained by the introduction of control interventions in the population in 2014, as demonstrated in previous studies^4,31^. The uneven distribution of sample size between the different age groups observed for both time points was due to limited availability of PRISM samples, particularly samples belonging to the “above18 years” age group with only one *Pf* positive sample for 2016.

**Table 2:** Summary statistics of study population sample size (n) and frequency (%), including malaria infection status according to microscopy and/or LAMP results, split by year of sample collection and stratified by age group. A large proportion of samples in 2013 are paired to those in 2016, graphically displayed in Supplementary Figure 2.

|  | 2013, n (%) | 2016, n (%) | Total, n (%) |
| --- | --- | --- | --- |
| **Sample size** | 310 (61.39) | 195 (38.61) | 505 (100.00) |
| **Age strata** |  |  |  |
| 6 months – 5 years | 71 (22.90) | 45 (23.08) | 116 (22.97) |
| 5 years – 11 years | 152 (49.03) | 139 (71.28) | 291 (57.62) |
| > 18 years | 87 (28.06) | 11 (5.64) | 98 (19.41) |
| **Malaria positive by age group** |  |  |  |
| 6 months – 5 years | 49 (15.81) | 6 (3.08) | 55 (10.89) |
| 5 years – 11 years | 116 (37.42) | 33 (16.92) | 149 (29.50) |
| > 18 years | 52 (16.77) | 1 (0.51) | 53 (10.50) |
| **Total** | 217 (70.00) | 40 (20.51) | 257 (50.89) |

### 3.3 Age-related seropositivity

There is an overall reduction in seroprevalence for the majority of the panel antigens tested, including the STEVOR and RIFIN recombinants, between 2013 and 2016, most prominently observed in the “6 months to 5 years” and “5 years to 11 years” age groups, as shown in Figure 3, except for the RIFIN3_SC recombinant with higher seroprevalence in 2016 samples compared to 2013. However, this trend was not reflected in the “above 18 years” age group, potentially confounded by the large samples size difference between the two time points with only one confirmed malaria positive sample in 2016, compared to 52 samples in 2013. For the 2013 samples (pre-intervention), seroprevalence for all short-term markers increased with age. In contrast, in 2016 (post-intervention), seroprevalence to Etramp5.Ag1, Etramp4.Ag2 and GEXP18 decreased with age. For all the long-term exposure markers, seroprevalence increased with age at both timepoints. Responses to MSP2.Dd2 in 2013 followed the same tendency but by contrast, in 2016, there was a decline of seroprevalence with increasing age, although not significant. Higher seroprevalence was observed against the tested recombinants semiconserved domains, especially for STEVOR1_SC, except for RIFIN3_SC, that presents a lower seroprevalence compared to the variable region. For the youngest age group there was little to no seroprevalence against all the variable domains of the recombinants, compared to their semiconserved regions, possibly reflecting a lack of exposure to the variant types used. Seropositivity thresholds are summarized in Supplementary Table 2, and absolute values to the relative percentage values are summarized in Supplementary Table 3.

**Figure3:**
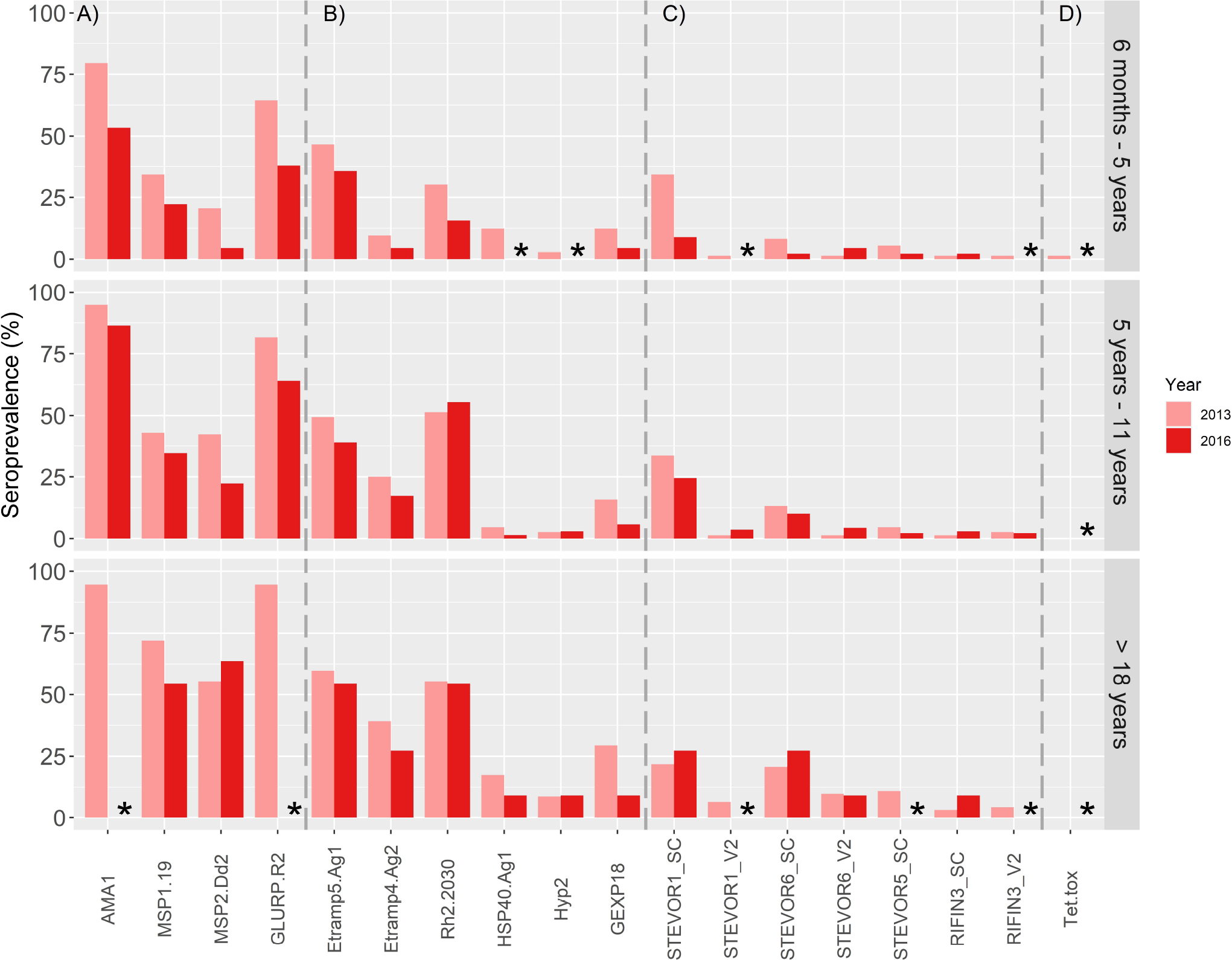
Population seroprevalence (%) per survey year for A) long-term markers; B) short, medium-term markers C) STEVOR/RIFIN recombinants and D) internal serology control, stratified by age group. Seroprevalence threshold was selected using a cut off mean MFI from PHE negative samples + 3 times standard deviation. Bar plots are paired colour coded, per recombinant, for visual representation of difference in seroprevalence between the two survey years. Tet.tox represents the internal positive human serological control. Seroprevalence of 0% is indicated with an Asterix and exact seroprevalence percentages are summarized in Supplementary Table 2.

The odds of being seropositive in 2016 as opposed in 2013 are lower in the “6 months to 5 years” age group for all antigens except for AMA1 and MSP2.Dd2 long-term markers OR=0.276 (95% CI 0.11-103.01; p=0.001) and OR=0.177 (95%CI 0.02-60.15; p=0.026) respectively, indicated in Figure 4A. For the adolescent group of 5 years to 11 years of age, the odds of seropositivity in 2016 compared to 2013 was greater for both short and long-term markers of malaria exposure (Figure 4B), where for the adults of above 18 years of age can be seen that the odds for most of the markers to be seropositive are also high in 2016, despite the interventions (Figure 4C). This is to be expected, since acquired immunity to malaria is age and exposure dependent and seropositivity in adults exposed to the infection for longer time periods is expected to be higher. For the tested STEVOR and RIFIN recombinants, odds of seropositivity also increased with age, but this was not as pronounced compared to the other groups of markers. It can be noted that the precision of the OR estimate for the STEVOR and RIFIN recombinants is higher overall, illustrated with larger squares and shorter confidence intervals. Antigens which have 0% seropositivity in one or both time points are not included in the OR analysis, as having 0 in the denominator will result in infinite odds. In the oldest age group, we can see an abnormality of 0% and almost 100% seropositivity juxtaposition for STEVOR1_SC recombinant between 2016 and 2013, respectively, due to the small sample size (n=11) in this age group for 2016.

**Figure 4:**
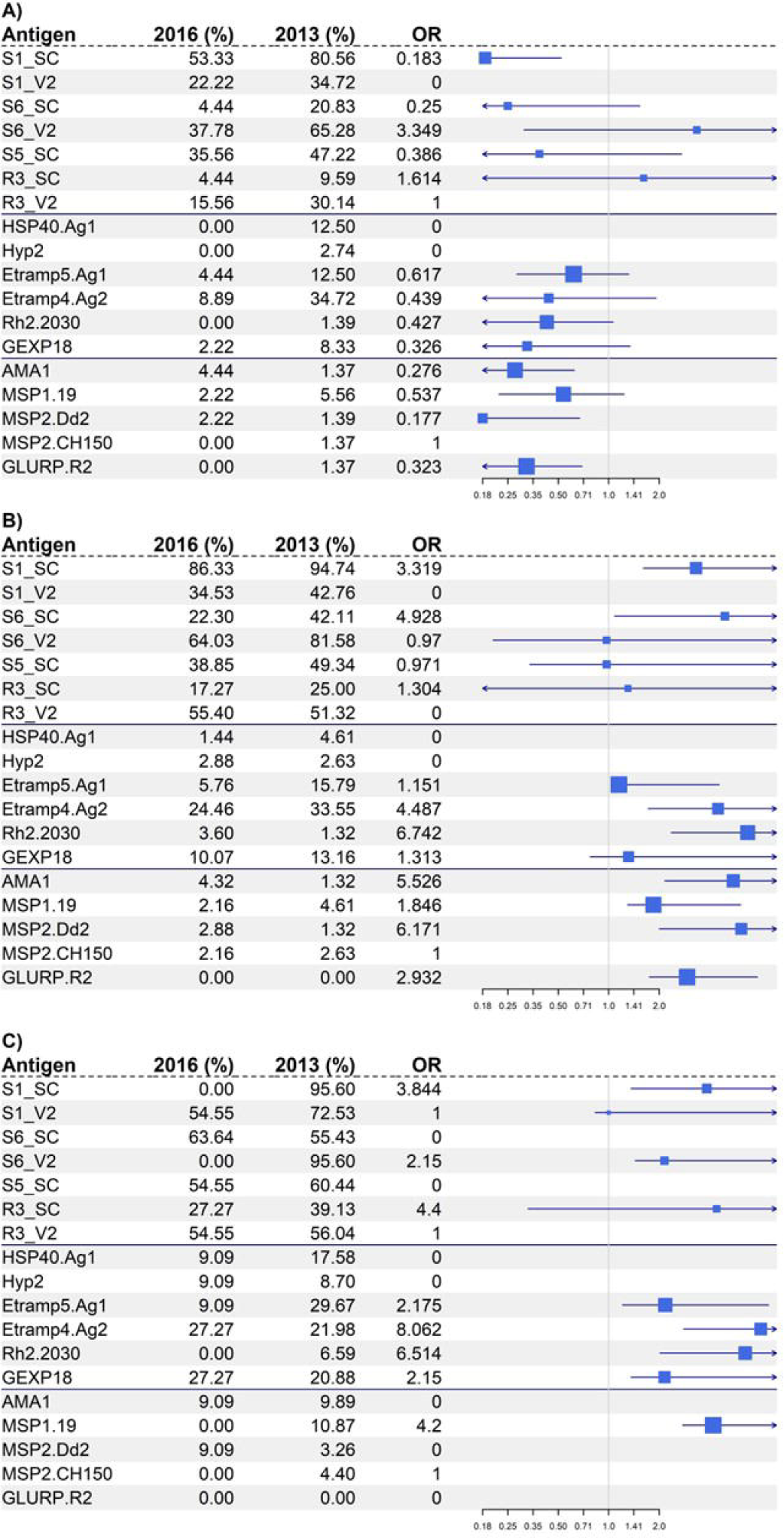
Forest plot of odds ratio analysis per recombinant antigen of being seropositive in 2016 compared to 2013, calculated in seroprevalence (%), stratified by age: A) 6 months to 5 years old, B) 5 years to 11 years old, and C) more than 18 years old. Squares graphically represent the calculated and summarized in the tables ORs with 95% confidence intervals (CIs) as whiskers. The size of the square is proportional to the level of precision of the calculated OR values (shorter CIs), the larger the square, the higher the precision. Recombinant antigens towards which there is 0% seropositivity, seen in the summary table on the left are not included in the OR analysis. ORs. of 10 or above of being seropositive in 2016 compared to 2013 are shown as 0 to omit them from the plot, due to large discrepancy of sample size resulting in abnormally high OR.

## 4. Discussion

This is the first study demonstrating methods for expression of STEVOR and RIFIN protein domains as recombinant proteins, exploring their potential use as serosurveillance tools by comparing them to previously confirmed markers of *P. falciparum* exposure, using the large multiplex suspension technology, Luminex ^16,26,32^.

In addition, this study compared two methods of recombinant expression in *E. coli* bacterial systems: a single GST tag plasmid and a dual His-Myc tag plasmid technology (Table 1). Higher reactivity measured as median fluorescence intensity (MFI) was observed for GST tagged recombinants on the Luminex multiplex serology platform displayed as protein titration plots in Supplementary Figure 1. This difference could be caused by the nature of the protein domains expressed as recombinants. Due to the protein design, the SC recombinants tend to express better with the single tag GST system, and because of their higher conservation between variants, it is expected higher serorecognition compared to the V2 recombinants^35^. The dual tag recombinants represent the V2 domains of selected STEVOR members which are expected to generate comparatively lower antibody response due to antigenic variation and the lack of knowledge if the samples tested in this study have been exposed to these specific variants. This trend is also observed in the ELISA results of the tested control pools, displayed in the last plot of the series in Figure 2. Variations in protein expression even within a single tag system is an established factor, clearly demonstrated in this study where the SC domain in the STEVORs expressed better as a GST tag system, where the GST system was preferred for the V2 domain of the studied RIFIN protein^36,37^. Secondary, some of the reactivity detected could potentially be against the GST tag molecule alone which is a naturally occurring enzyme molecule in the parasite *Schistosoma japonicum*^38^. There was no data available about past infections of *Schistosoma* for the participants used in this study, hence it was impossible to determine if there was natural antibody reactivity to the GST tag. To eliminate this as a possibility, a GST recombinant alone was included in the recombinant antigen panel used against the tested population for potentially subtracting reactivity values detected against the tag alone. However, a decision not to subtract the GST seroreactivity from the detected MFI values against the GST tagged recombinants was made, as after performing the seropositivity threshold analysis there were no antibody responses against GST detected above the threshold for any of the samples, summarized in Supplementary Table 2. Moreover, GST tag is used as a solubility factor which perhaps could influence the performance of the recombinants ^34^. This argument is supported by the detected precipitation of the His-Myc dual tag recombinants. Nonetheless, there was no significant difference in protein yield when comparing both systems, thus a decision that the tag expression system did not significantly affect the production of recombinants was made and the recombinants with the system resulting in higher yield was individually selected per recombinant. The fact that two different protein expression tag systems were used for this study and the recombinants used differ between each other can be regarded as a limitation in this study. However, as these were not confirmational proteins, there was little need to explore other more disparate expression systems such as yeast or wheat germ. The proposed construct design was based on expertise within the laboratory group and lent itself well to expression of the targets irrespective of the tag system used and, moving forward the optimum solution will be to focus on one expression system only ^39,40^.

The kinetics and half-life of antibodies to specific *Pf* antigens varies depending on multiple factors, such as protein antigenic properties, human host age and genetics, seasonality, immunoglobulin class, and endemicity of the setting ^14,15^. The antibody responses to certain antigens have a short half-life and thus indicate recent exposure to infection (short-term markers), such as HSP40.Ag1 (Heat Shock Protein 40 Antigen 1), Hyp2 (Hypothetical protein 2), Etramp5.Ag1 (Early Transcribed Membrane Protein 5 Antigen 1), Etramp4.Ag2 (Early Transcribed Membrane Protein 4 Antigen 2) and GEXP18 (Gametocyte Export Protein 18). Some already established markers, such as Rh2.2030 (Reticulocyte-binding protein homologue) and EBAs (Erythrocyte-binding antigens), indicate *Pf* infection during the past 6 months and therefore are considered as moderate-exposure markers ^16^. Antibody responses to AMA1 (Apical Membrane Antigen 1), MSP1.19 (Merozoite Surface Protein 1), MSP2.Dd2 (Merozoite Surface Protein 2, Dd2 allele), and GLURP.R2 (Glutamate Rich Protein Region 2) indicate longer-term exposure and can persist for years after infection, termed as long-term markers ^16,26,32^. The measurement of antibody levels against different types of immune markers is a useful tool to assess the time of infection in individuals or in a population. This level of information is of a high importance for the design of control programs to interrupt transmission in a specific setting ^26,41^. In concordance with other molecular techniques, measurement of antibody levels can also inform about the immune profiles in individuals protected from severe malaria disease, essential information for new vaccine approaches ^26,42^. High throughput multiplex platforms, like quantitative suspension array technologies (qSAT) used in this study, have been developed to allow sensitive and accurate estimates of time of exposure via serosurveillance, testing antibody titers against multiple targets simultaneously ^29^.

Tested samples selected from the two time points of the cohort, 2013 and 2016, were predominantly paired samples, with 74% of the 2016 samples paired with those from 2013. Thus, observing differences in antibody responses to the recombinants before and after intervention is done on the same population, where some of the samples would move from one age group to another between 2013 and 2016. This sample selection allows for an evaluation of the reduction in seroreactivity and serorecognition over time due to the implementation of interventions, leading to a reduction in *P. falciparum* malaria prevalence and incidence in 2016. It is also important to highlight that, after implementing interventions, there is an expected decrease of naturally acquired immunity (NAI) to *P. falciparum* due to decreased exposure, clearly illustrated in the seroprevalence plot in Figure 3 ^43^. However, the seropositivity analysis shows higher population seroprevalence to a few of the variant recombinants (STEVOR1_SC, STEVOR6_SC, RIFIN3_SC) in 2016 as opposed to 2013 for the “above 18 years” age group. There are a few possible explanations for this observation. These recombinants represent the semi-conserved regions of the variants to which higher serorecognition is expected due to high domain conservation, leading to cross reactivity between variants^35^. In addtion, the low sample size (n=11) of the adult age group in 2018 can significantly impact the calculation of seroprevalence and introduce bias to the analysis and interpretation of the data ^44^. Consequently, the focus of further discussion is directed mainly on the results from the other two age groups, which in turn is with a higher importance to the posed question since particularly children under 5 years of age have a naïve immune system and changes in antibody responses due to external or internal pressure can be detected with higher precision and sensitivity, as well as the adolescent group is known to be the age group with the most exposure to infection ^45^. Another potential limitation was the fact that despite the selected high reactivity serum as a control, it is still possible that some of the recombinants reactivity is not captured due to their hypervariability nature.

While few studies have been conducted analyzing the humoral response of STEVORs and RIFINs but, similar to *Pf*EMP1, IgG antibody responses have been correlated with age and it has been observed that children with high anti-RIFIN and anti-STEVOR antibody titers had a reduced risk of febrile malaria ^46^. Consistent with these findings, there is an observed increase of seroprevalence to STEVORs and RIFINs with age (Figure 3), most pronounced in the case of STEVOR6_SC. Comparing calculated seroprevalence between the RIFINs and STEVORs, there is an overall higher seropositivity for the latter, in contrast to a study performed in Uganda that showed a higher seroprevalence against RIFIN compared to STEVOR variants^46^. The difference of findings could be explained by the presence of only one RIFIN variant, as a recombinant protein in this study, as compared to around 150 *rif* peptides analyzed in the study by Kanoi B *et al*. Moreover, some 3D7 strain RIFINs have been found to bind leukocyte immunoglobulin-like receptor B1 (LILRB1) or leucocyte-associated immunoglobulin-like receptor 1 (LAIR1), suppressing immune responses and inhibiting natural killer (NK) effector cell activation ^47^.

Furthermore, all age groups have some seroprevalence calculated against the semi conserved recombinants, where for the variable domain recombinants the younger age groups present with 0% seroprevalence, perhaps reflecting the lack of exposure to these variants. The fact that infants lose their maternal antibodies after 6 months of age could explain these results, as the variable domain forms a surface-exposed loop that is highly variable due to a high recombination rate, a mechanism that the parasite uses to create variability in the proteins, for immune system evasion, compared to the semiconserved region ^13,48^. Additionally, their location in the subtelomeric region of the parasite’ s genome facilitates the mechanisms that induce hypervariability within genes, playing a role in *P. falciparum’ s* antigenic variation^13,49^. Apart from their high variability, they are under the host’ s immune pressure which amplifies the variable repertoire of these proteins ^49,50^. Since the SC domain has higher likelihood of recognition compared to V2, it is possible that contribution to the acquisition of NAI is restricted to the semiconserved region. Further studies comprised of a larger library of variants from the STEVOR and RIFIN protein families are needed to test this hypothesis. The odds ratio (OR) analysis, summarized and graphically represented in Figure 4, shows odds of increased seropositivity in 2016 compared to 2013 is greater in the older age groups. Moreover, in the older age groups as compared to the infants, there was a greater magnitude of increased odds of seropositivity to long-term markers, compared to short and medium-term markers, and the STEVOR and RIFIN recombinants. These observations are all expected due to the known age and exposure dependent acquired immunity to *P. falciparum* exposure ^14,16^. Nevertheless, all markers of exposure as well as the tested novelle recombinants show higher odds of seropositivity with increasing age, visualized in Figure 4.

This study reports the first STEVOR and RIFIN variants expressed as antigen recombinants using various *in silico* tools and an *E. coli* bacterial expression system. Moreover, since the tested STEVOR and RIFIN recombinants follow the same trends of seropositivity to the already established panel of antigen markers of exposure, it can be hypothesized that members of those protein families could potentially be used as novel markers of *P. falciparum* exposure in serosurveillance ^16,26^. However, due to the hypervariability between variants, further studies comprised of a larger library of variants representing the STEVOR and RIFIN families should be compiled to strengthen the findings of this study.

## 9. List of non-standard abbreviations

*Pf*: *Plasmodium falciparum*
IE: Infected Erythrocyte
VSA: Variant Surface Antigen
*Pf*EMP1: *Plasmodium falciparum* Erythrocyte Membrane Protein 1
RIFIN: Repetitive Interspersed Family
STEVOR: Subtelomeric Variable Open Reading Frame family
*Pf*MC-2TM: *Plasmodium falciparum* Maurer’ s Cleft 2 Transmembrane family
SURFIN: Surface Associated Interspersed family.
SP: Signal Peptide
SC: Semi Conserved domain
V1, V2: Variable domain 1, Variable domain 2
HSP40.Ag1: Heat-Shock Protein 40
Hyp2: Hypothetical Protein 2
Etramp5.Ag1: Early Transcribed Membrane Protein 5
Etramp4.Ag2: Early Transcribed Membrane Protein 4
GEXP18: Gametocyte Export Protein 18
Rh2.2030: Reticulocyte Binding Protein Homologue 2
EBAs: Erythrocyte Binding Antigens
AMA1: Apical Membrane Antigen 1
MSP1.19: Merozoite Surface Antigen 1
MSP2.Dd2: Merozoite Surface Antigen 2, Dd2 allele
MSP2.CH150: Merozoite Surface Antigen 2, CH150 allele
GLURP.R2: Glutamate-Rich Protein, Region 2
PRISM: Program for Resistance, Immunology, Surveillance and Modeling of Malaria in Uganda
DNTMaPa: Malaria and Neglected Tropical Diseases mapping survey, Guinea-Bissau.
GST: Glutathione S-transferase protein
LAMP: Loop-mediated Isothermal Amplification
MFI: Median Fluorescence Intensity
ITN: Insecticides Treated Nets
IRS: Indoor Residual Spraying
OR: Odds Ratio
CI: Confidence Interval
LILRB1: Leucocyte Immunoglobulin-Like Receptor B1
LAIR1: Leucocyte-associated Immunoglobulin Receptor 1
NK: Natural Killer cell

## 10. Acknowledgments

Special thanks to Dr Christian Kositz from Swiss TPH for analytical advice, as well as the data science department in the MRC Unit The Gambia @ LSHTM.

