## Supplementary materials file for "Novel hypervariable erythrocyte surface expressed recombinant proteins show promise as serological markers of exposure to *Plasmodium falciparum* infection"

### 1 Supplementary Figures

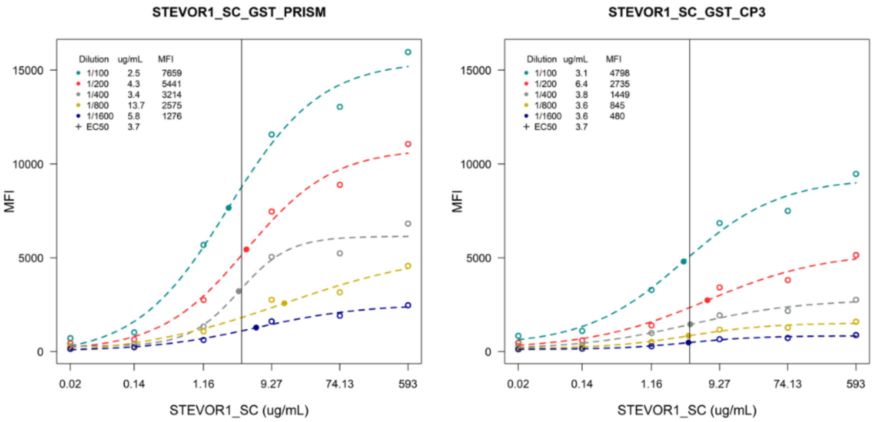

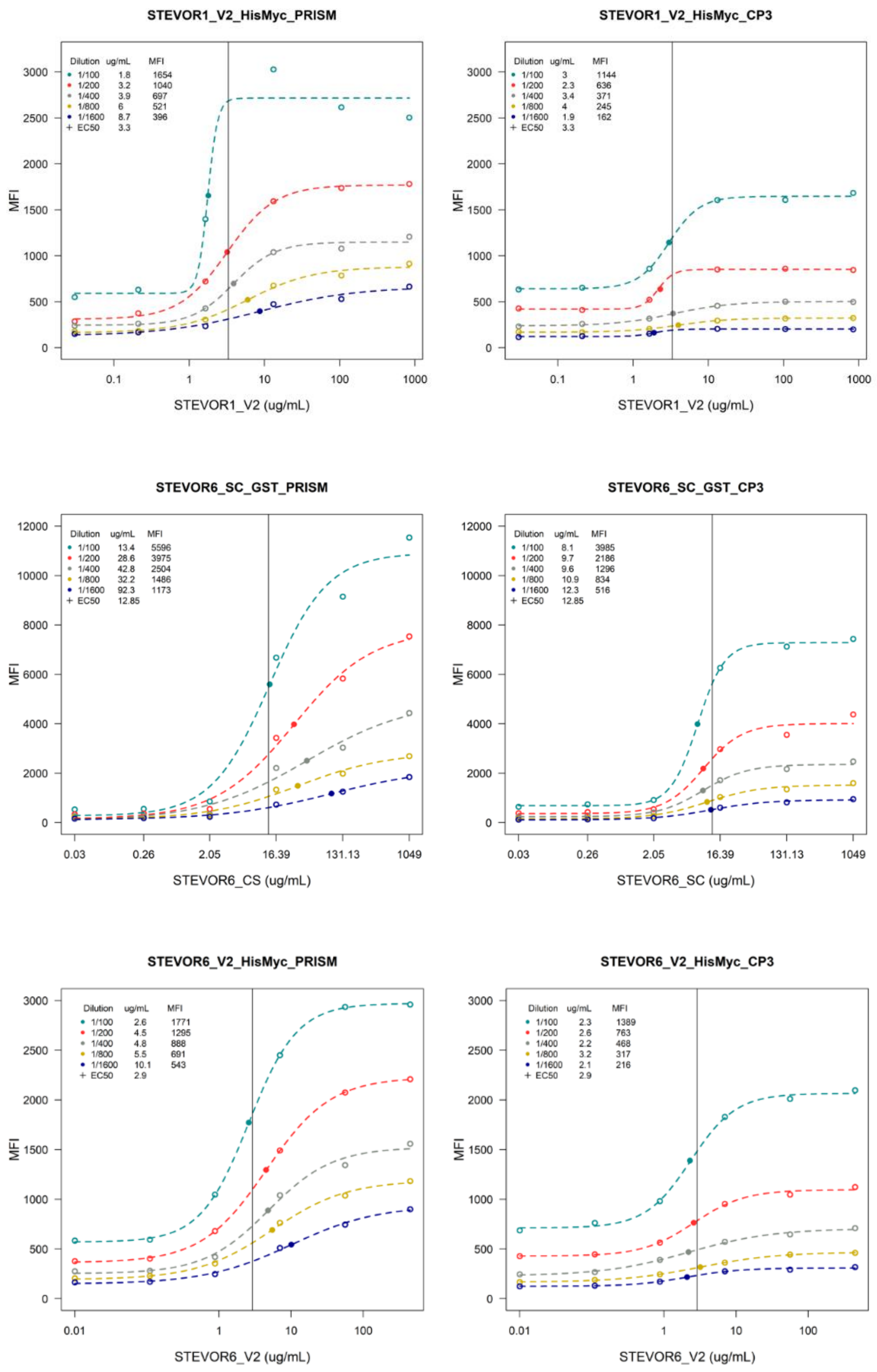
STEVOR1_V2_HisMyc_PRISM
STEVOR1_V2_HisMyc_CP3
MFI
STEVOR1_V2 (ug/mL)
STEVOR6_SC_GST_PRISM
STEVOR6_SC_GST_CP3
STEVOR6_CS (ug/mL)
STEVOR6_SC (ug/mL)
STEVOR6_V2_HisMyc_PRISM
STEVOR6_V2_HisMyc_CP3
STEVOR6_V2 (ug/mL)

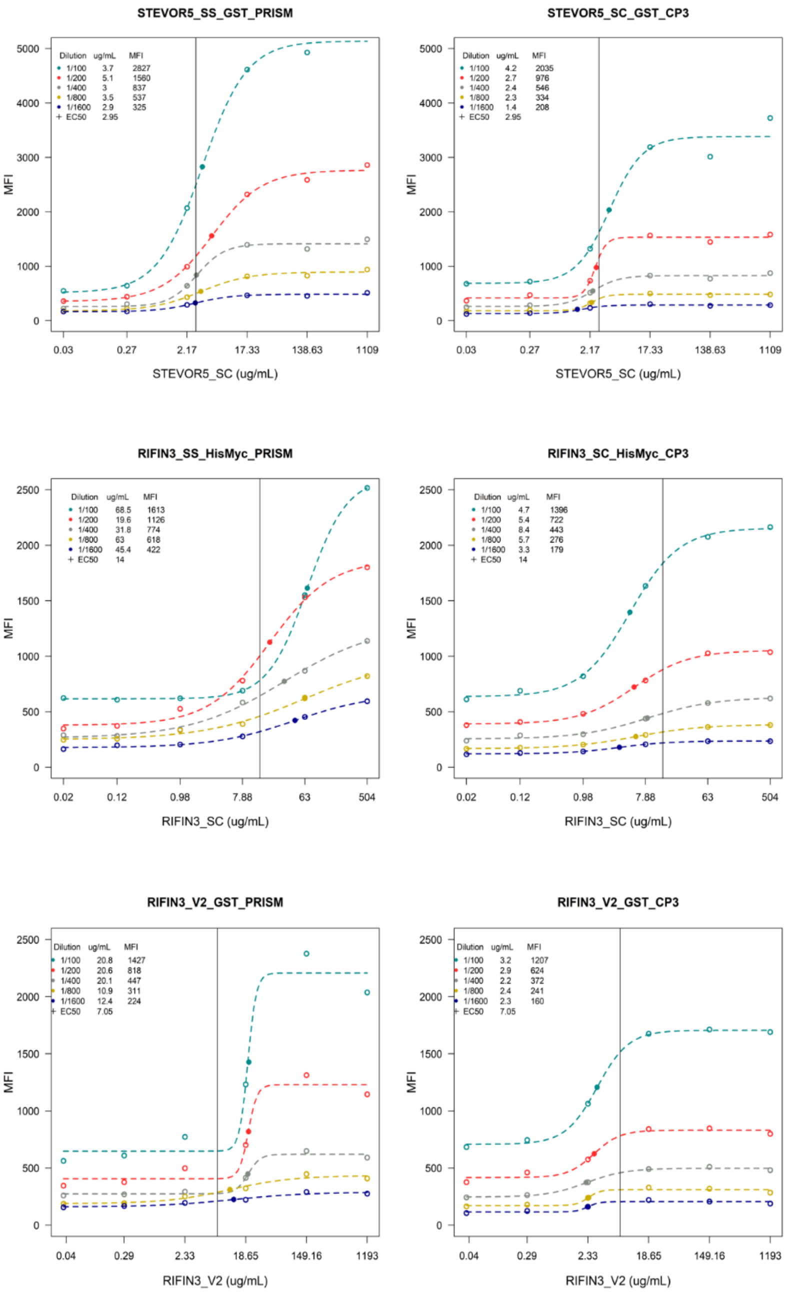


**<u>Supplementary Figure 1:</u> Recombinant proteins titrations on MagPlex microsphere beads against two positive controls pool serum from Tanzania (CP3) and Uganda (PRISM1). Optimum coupling concentration (EC50 point of saturation) was calculated using the medium concentration value from all curves subjected to 4-parameter logistic regression.**

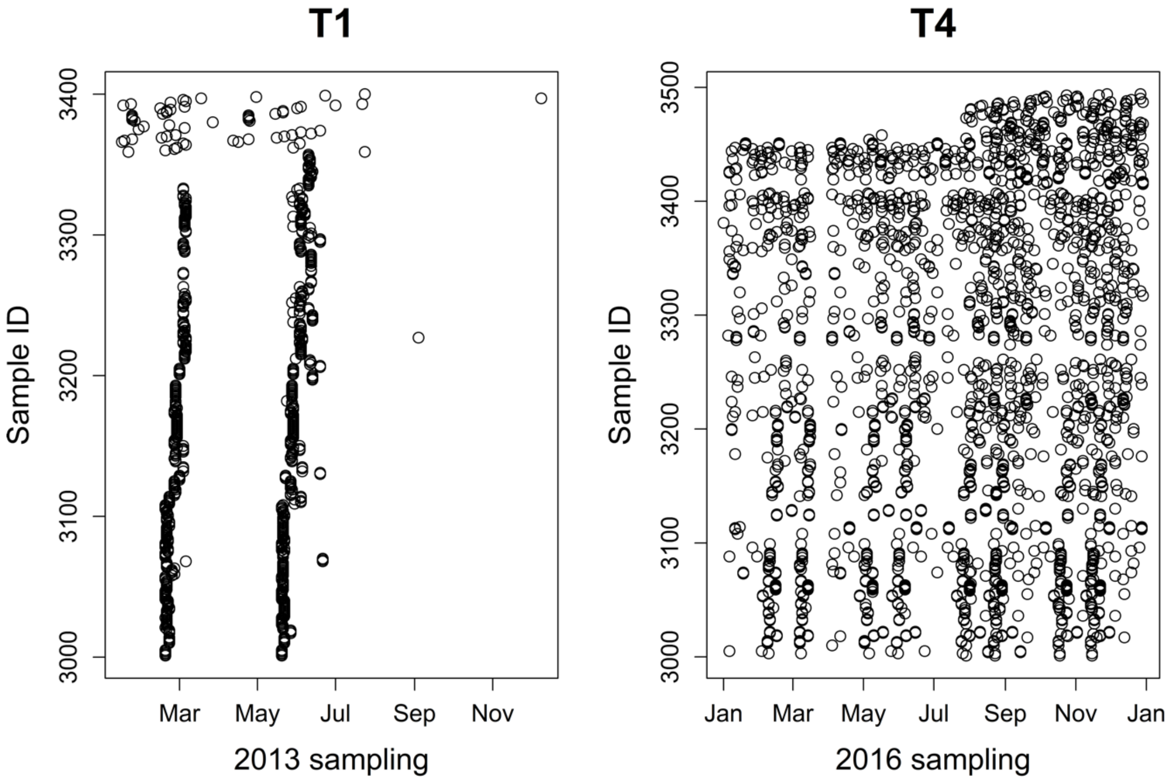


**<u>Supplementary Figure 2:</u> Sampling distribution time of the year for the two sample time points (2013 and 2016). Sample IDs are displayed on the y-axis, indicating that the majority of the samples are paired.**

### 2 Supplementary tables

**<u>Supplementary Table 1:</u> Summary of already established *Plasmodium falciparum* markers of exposure complementing the studied STEVOR and RIFIN recombinant proteins.**

| **Recombinant Antigen** | **Protein Name** | **Protein Species** | **Exposure marker** | **Expression system** |
|---|---|---|---|---|
| AMA1 | Apical Membrane Antigen 1 | *Plasmodium falciparum* | Long-term marker | His-tag *E. coli* |
| MSP1.19 | Merozoite Surface Protein 1 | *Plasmodium falciparum* | Long-term marker | GST-tag *E. coli* |
| GLURP.R2 | Glutamate-Rich Protein Region 2 | *Plasmodium falciparum* | Long-term marker | GST-tag *E. coli* |

| | | | | |
|---|---|---|---|---|
| MSP2.Dd2 | Merozoite Surface Protein 2, Dd2 allele | *Plasmodium falciparum* | Short-term marker | GST-tag *E. coli* |
| Etramp5.Ag1 | Early Transcribed Membrane Protein 5 | *Plasmodium falciparum* | Short-term marker | GST-tag *E. coli* |
| Etramp4.Ag2 | Early Transcribed Membrane Protein 4 | *Plasmodium falciparum* | Short-term marker | GST-tag *E. coli* |
| Rh2.2030 | Reticulocyte Binding Protein Homologue 2 | *Plasmodium falciparum* | Medium-term marker | GST-tag *E. coli* |
| HSP40.Ag1 | Heat-Shock Protein 40 | *Plasmodium falciparum* | Short-term marker | GST-tag *E. coli* |
| Hyp2 | Hypothetical Protein 2 | *Plasmodium falciparum* | Short-term marker | GST-tag *E. coli* |
| GEXP18 | Gametocyte Exported Protein 18 | *Plasmodium falciparum* | Short-term marker | GST-tag *E. coli* |
| Tet.tox | Tetanus toxoid | *Clostridium tetani* | Immunization antigen, serology control | GST-tag *E. coli* |
| GST | Glutathione-S-Transferase | *Schistosoma japonicum* | GST-tagged proteins control | GTS-tag *E. coli* |

**<u>Supplementary Table 2:</u> Summary table of mean MFI values of PHE negative controls and calculated seropositivity thresholds of PHE mean MFI plus three times standard deviation, per antigen.**

| Antigen | Mean PHE (MFI) | 3SD cut-off (MFI) |
|---|---|---|
| STEVOR1_SC | 303.20 | 1580.84 |
| STEVOR1_V2 | 146.43 | 463.57 |
| STEVOR6_SC | 253.65 | 1182.37 |
| STEVOR6_V2 | 229.97 | 692.81 |
| STEVOR5_SC | 221.48 | 651.31 |
| RIFIN3_SC | 220.61 | 1044.30 |
| RIFIN3_V2 | 245.83 | 767.67 |
| HSP40.Ag1 | 962.02 | 2861.01 |
| Hyp2 | 894.38 | 2595.58 |
| Etramp5.Ag1 | 260.74 | 725.93 |
| Etramp4.Ag2 | 979.62 | 3044.99 |
| Rh2.2030 | 378.96 | 1609.28 |
| GEXP18 | 518.18 | 1712.87 |
| AMA1 | 220.38 | 662.60 |
| MSP1.19 | 792.07 | 2362.44 |
| MSP2.Dd2 | 1022.31 | 11100.98 |

| | | |
|---|---|---|
| GLURP.R2 | 153.93 | 478.74 |
| Tet.tox | 13019.20 | 26842.07 |
| GST | 337.34 | 985.23 |

**Supplementary Table 3: Summary of population seroprevalence per survey year for each recombinant antigen.**

| | 2013 Seroprevalence n (%) | | | 2016 Seroprevalence n (%) | | |
|---|---|---|---|---|---|---|
| **Age strata** | **6 months – 5 years** | **5 years – 11 years** | **>18 years** | **6 months – 5 years** | **5 years – 11 years** | **>18 years** |
| **Long-term markers** | | | | | | |
| PfAMA1 | 58 (79.45) | 144 (94.74) | 87 (94.57) | 24 (53.33) | 120 (86.33) | 0 (0.00) |
| PFMSP1.19 | 25 (34.25) | 65 (42.76) | 66 (71.74) | 10 (22.22) | 48 (34.53) | 6 (54.55) |
| MSP2.Dd2 | 15 (20.55) | 64 (42.11) | 51 (55.43) | 2 (4.44) | 31 (22.30) | 7 (63.64) |
| GLURP.R2 | 47 (64.38) | 124 (81.58) | 87 (94.57) | 17 (37.78) | 89 (64.03) | 0 (0.00) |
| **Short-term markers** | | | | | | |
| HSP40.Ag1 | 9 (12.33) | 7 (4.61) | 16 (17.39) | 0 (0.00) | 2 (1.44) | 1 (9.09) |
| Hyp2 | 2 (2.74) | 4 (2.63) | 8 (8.70) | 0 (0.00) | 4 (2.88) | 1 (0.00) |
| Etramp5.Ag1 | 34 (46.58) | 75 (49.34) | 55 (59.78) | 16 (35.56) | 54 (38.85) | 6 (54.55) |
| Etramp4.Ag2 | 7 (9.59) | 38 (25.00) | 36 (39.13) | 2 (4.44) | 24 (17.27) | 3 (27.27) |
| Rh2.2030 | 22 (30.14) | 78 (51.32) | 51 (55.43) | 7 (15.56) | 77 (55.40) | 6 (54.55) |
| GEXP18 | 9 (12.33) | 24 (15.79) | 27 (29.35) | 2 (4.44) | 8 (5.76) | 1 (9.09) |
| **Hypervariable recombinants** | | | | | | |

| | | | | | | |
|---|---|---|---|---|---|---|
| STEVOR1_SC | 25 (34.25) | 51 (33.55) | 20 (21.74) | 4 (8.89) | 34 (24.46) | 3 (27.27) |
| STEVOR1_V2 | 1 (1.37) | 2 (1.32) | 6 (6.52) | 0 (0.00) | 5 (3.6) | 0 (0.00) |
| STEVOR6_SC | 6 (8.22) | 20 (13.16) | 19 (20.65) | 1 (2.22) | 14 (10.07) | 3 (27.27) |
| STEVOR6_V2 | 1 (1.37) | 2 (1.32) | 9 (9.78) | 2 (4.44) | 6 (4.32) | 1 (9.09) |
| STEVOR5_SC | 4 (5.48) | 7 (4.61) | 10 (10.87) | 1 (2.22) | 3 (2.16) | 0 (0.00) |
| RIFIN3_SC | 1 (1.37) | 2 (1.32) | 3 (3.26) | 1 (2.22) | 4 (2.88) | 1 (9.09) |
| RIFIN3_V2 | 1 (1.37) | 4 (2.63) | 4 (4.35) | 0 (0.00) | 3 (2.16) | 0 (0.00) |
| **Internal Control** | | | | | | |
| Tet.tox | 1 (1.37) | 0 (0.00) | 0 (0.00) | 0 (0.00) | 0 (0.00) | 0 (0.00) |
| GST | 0 (0.00) | 0 (0.00) | 0 (0.00) | 0 (0.00) | 0 (0.00) | 0 (0.00) |
